# Identification and Characterization of Adult Islet Pancridia Cells Capable of Differentiating into Islet Organoids

**DOI:** 10.1101/2025.07.28.667223

**Authors:** Carly M. Darden, Jayachandra Kuncha, Jeffrey T. Kirkland, Jordan Mattke, Srividya Vasu, Bashoo Naziruddin, Michael C. Lawrence

**Author notes:** Corresponding author: Michael C. Lawrence. These authors contributed equally (Carly M. Darden, Jayachandra Kuncha).

## Abstract

Multipotent progenitor-like cells have been identified in the adult pancreas under various physiological and pathological conditions. Here, we identify and characterize a subset of adult pancreas-derived cells, termed islet pancridia cells (IPCs), that can be expanded in vitro and retain the potential for endocrine differentiation and islet cell function. Single-cell RNA sequencing of expanded pancridia revealed transcriptomic profiles resembling immature beta cells, enriched with markers of epithelial-mesenchymal transition. A CD9⁺, PROCR⁺ subpopulation of IPCs formed IPC clusters marked by restricted expression of RGS16, a known islet progenitor marker. In vivo, co-transplantation of expanded IPCs with a subtherapeutic dose of islets significantly improved graft function and partially restored native pancreatic endocrine activity in streptozotocin-induced diabetic mice. In vitro, treatment of RGS16⁺ IPC clusters with the small molecule ISX9 induced differentiation into islet organoids that co-expressed and secreted insulin and glucagon. ISX9-mediated differentiation was driven by calcineurin/NFAT-dependent recruitment of the histone acetyltransferase p300 and displacement of histone deacetylases (HDACs) at the RFX6 and NEUROD1 promoters. Pre-treatment with the HDAC inhibitor ITF2357 further enhanced islet cell differentiation by promoting chromatin remodeling and facilitating NFAT-targeted recruitment of p300. These findings uncover calcium-dependent and epigenetic mechanisms that regulate the differentiation of multipotent CD9⁺, PROCR⁺, RGS16⁺ IPCs into functional islet organoids and offer potential strategies for regenerating islet cell mass to treat diabetes.

## INTRODUCTION

Islet transplantation offers a promising cell replacement therapy to prevent or reverse diabetes in individuals who have lost functional beta and alpha cells. However, the limited availability of donor islet tissue remains a major barrier to its broader clinical application. This limitation has driven intense investigation into alternative strategies, including in vitro islet cell engineering and beta-cell expansion to restore or maintain islet cell mass and function.

Significant progress has been made in generating stem cell-derived “beta-like” cells by directing their differentiation through stepwise protocols that mimic developmental processes. These protocols guide human embryonic stem cells (hESCs) or induced pluripotent stem cells (iPSCs) through definitive endoderm and pancreatic progenitor stages to ultimately produce glucose-responsive, insulin-secreting cells (1–3). Most strategies employ 4 to 7 sequential culture stages that activate key transcription factors critical to islet development, including *PDX1*, *NEUROG3*, *RFX6*, *NEUROD1*, *NKX2-2*, *NKX6-1*, *MAFA*, and *MAFB*. (4–14).

Stem cell-derived islets (sc-islets) have demonstrated the ability to reverse diabetes in animal models and are currently being evaluated in phase I and II clinical trials (15). While their therapeutic potential is substantial, use of hESCs and iPSCs for treating type 1 diabetes (T1D) requires immunosuppression and raises concerns about tumorigenicity, oncogenic mutations, and ethical issues related to embryonic sources (16–18). As a result, alternative approaches using non-embryonic, non-genetically modified adult-derived cells, including chemically induced iPSCs, are under active investigation (19–23).

Studies have shown that insulin- and glucagon-producing cells can be derived from pancreatic ductal epithelial cells and mesenchymal stromal/stem cells (MSCs) (24–36). However, these source populations are highly heterogeneous, and the specific cell subtypes capable of endocrine differentiation remain poorly defined. Even cultures purified for canonical MSC surface markers contain diverse subpopulations with varying differentiation potentials, influenced by both tissue origin and culture conditions. Moreover, it remains unclear whether these cells can directly give rise to insulin-producing cells or primarily support the function and proliferation of existing beta cells.

A longstanding question in the field is whether an adult human stem cell population exists that can generate functional, glucose-responsive beta cells. Some studies have provided evidence of de novo beta-cell regeneration in both animal and human models (37–39). Yet, other lineage tracing studies in mice have demonstrated that beta cells are no longer produced once existing insulin-producing cells are ablated (40). These findings have led to alternative hypotheses, including the possibility that beta-cell regeneration arises not from pluripotent precursors, but from dedifferentiated or partially differentiated beta cells retained in a progenitor-like state (41–46).

Supporting this idea, we have observed that stressed or cultured beta cells can undergo dedifferentiation, and under specific conditions, regain insulin production and secretory function in vitro (47). Based on these observations, we hypothesized that beta cells can revert to a progenitor state, enabling their expansion and redifferentiation into functional beta-like cells. In this study, we establish a method to recover a population of adult pancreatic progenitors—termed islet pancridia cells—from islet isolation procedures. We provide proof of concept for their expansion and subsequent regeneration into insulin- and glucagon-releasing islet organoids in vitro. With further scale-up and optimization, this approach may enable production of autologous tissue for islet replacement therapies. Moreover, the signaling mechanisms identified in this study may offer therapeutic targets to regenerate islet organoids *in vivo* and restore endocrine function in diabetes.

## RESULTS

### Single Cell RNA-Seq Analysis of Expanded Pancreatic Tissue Reveals a Subpopulation of Islet Pancridia Cells in a Differentiated State

We used transcriptomic profiling to analyze single cells from cultured human pancreatic tissue to identify and characterize progenitor cells expanded in culture. Adult pancreatic samples were collected from COBE 2991 cell processor fractions during islet isolation procedures and expanded in culture for 8–15 passages (Suppl. Table 1). Single-cell RNA sequencing was then performed on dissociated cells to profile the expanded populations. A total of 100,000 cell barcodes were processed and analyzed using reference-based mapping against integrated human pancreatic tissue scRNA-seq datasets from six independent studies using various single-cell technologies compiled in the Human BioMolecular Atlas Program (HuBMAP) (48, 49).

Automated cell type annotation using Azimuth identified multiple pancreatic cell subtypes with transcriptomic signatures matching activated stellate cells, ductal cells, and beta cells (Fig. 1A) (50). Seurat analysis of highly variable genes and dimensionality reduction via t-distributed stochastic neighbor embedding (t-SNE) revealed 20 clusters with distinct expression profiles. A specific subpopulation of 16,381 cells spanning clusters 9, 10, 14, 17, and 18 co-expressed genes associated with islet cell progenitors and markers characteristic of ductal epithelium, stellate cells, and MSCs (Figs 1B, 1C, 1D; Suppl. Table 2).

**FIGURE 1.**
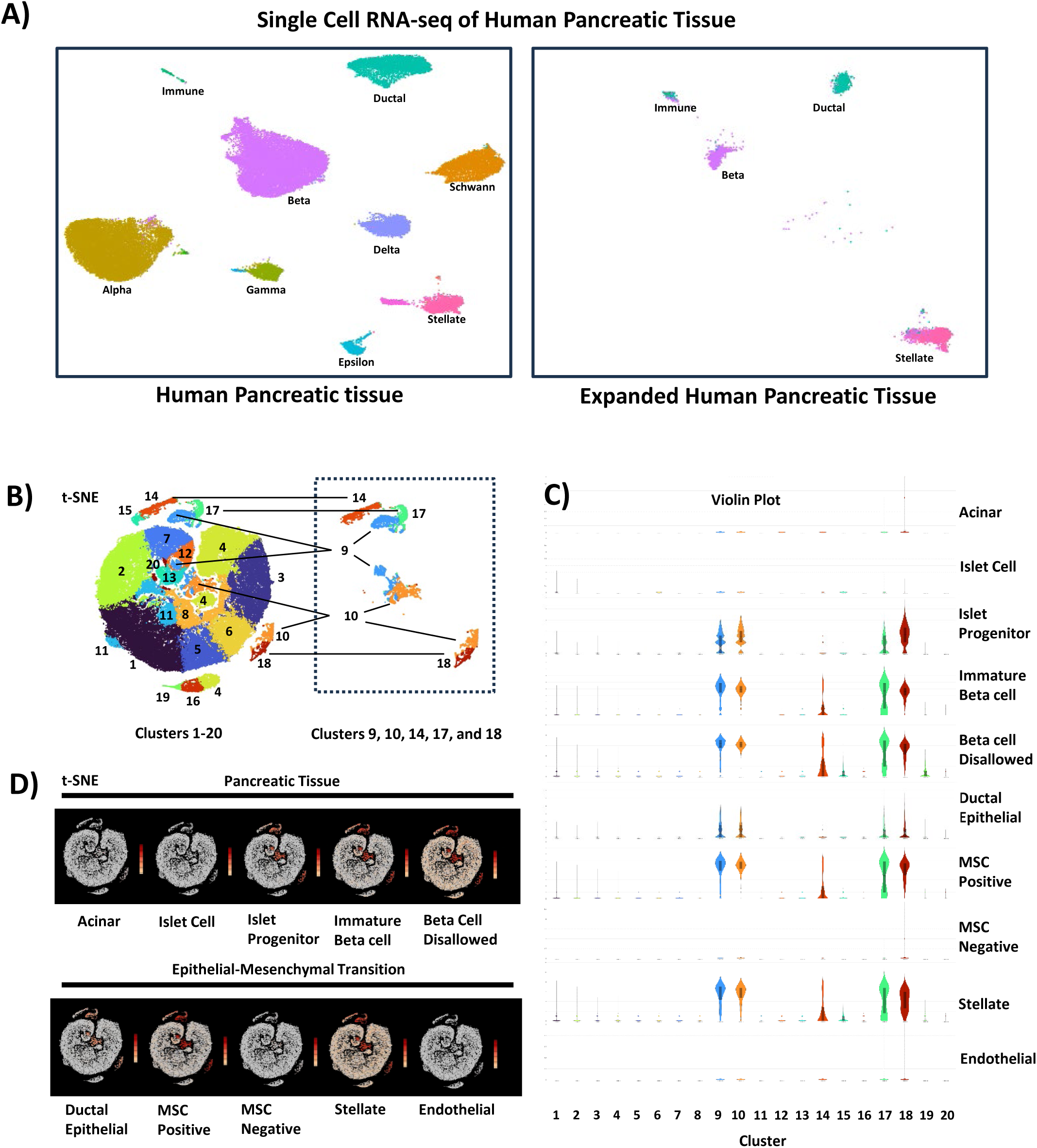
Identification of IPCs in expanded human pancreatic tissue by single cell transcriptomic profiling. (A) Azimuth automated cell type annotation mapping of scRNA-seq data from expanded human pancreatic tissue (100,000 cells) referenced to dataset aggregates (n=6) from human pancreatic tissue (35,000 cells) and (B) Seurat analysis of highly variant genes and dimensionality reduction by t-distributed stochastic neighbor embedding (t-SNE). (C) Gene expression profiling and (D) t-SNE mapping of cluster 9, 10, 14, 17, and 18 subpopulations containing islet cell progenitor cell markers (9,289 cells). Data shown are results from one donor.

Importantly, this subpopulation lacked expression of endothelial, acinar, or mature islet cell markers but expressed immature beta cell markers and “beta cell–disallowed” genes. These transcriptomic patterns suggest the expanded cells share features with dedifferentiated beta-like cells. We herein designate this subpopulation as islet pancridia cells (IPCs) to distinguish them from traditional pancreatic progenitor cells described during development or derived from embryonic stem cells.

### IPCs Express Multiple Progenitor Markers and Signaling Pathways Associated with Regeneration

Further reclustering and differential gene expression analysis of the IPC population revealed strong upregulation (>50-fold) of transcripts including *CD81*, *MMP2*, *TEAD1*, *YAP1*, *CD9*, *KRT19*, *BMPR1A*, *PPP3R1*, *PPP3CC*, *ALDH1A*, *ALDH1B1*, *HDAC1*, *HDAC3*, and *FOXO1* (Fig. 2A). Gene set enrichment analysis (GSEA) of the top 100 upregulated genes showed highest enrichment in pancreatic progenitor cell types, matched against the PanglaoDB and CellMarker 2.0 databases (Suppl. Table 3) (51, 52). The most enriched signaling pathways included those associated with epithelial-mesenchymal transition (EMT) and beta-cell development, based on matches to the MSigDB Human Molecular Signatures database (Suppl. Table 4A) (53). Gene Ontology analysis of Molecular Function revealed strong enrichment in *ALDH* activity and *BMP receptor* signaling (Suppl. Table 4B) (54, 55).

**FIGURE 2.**
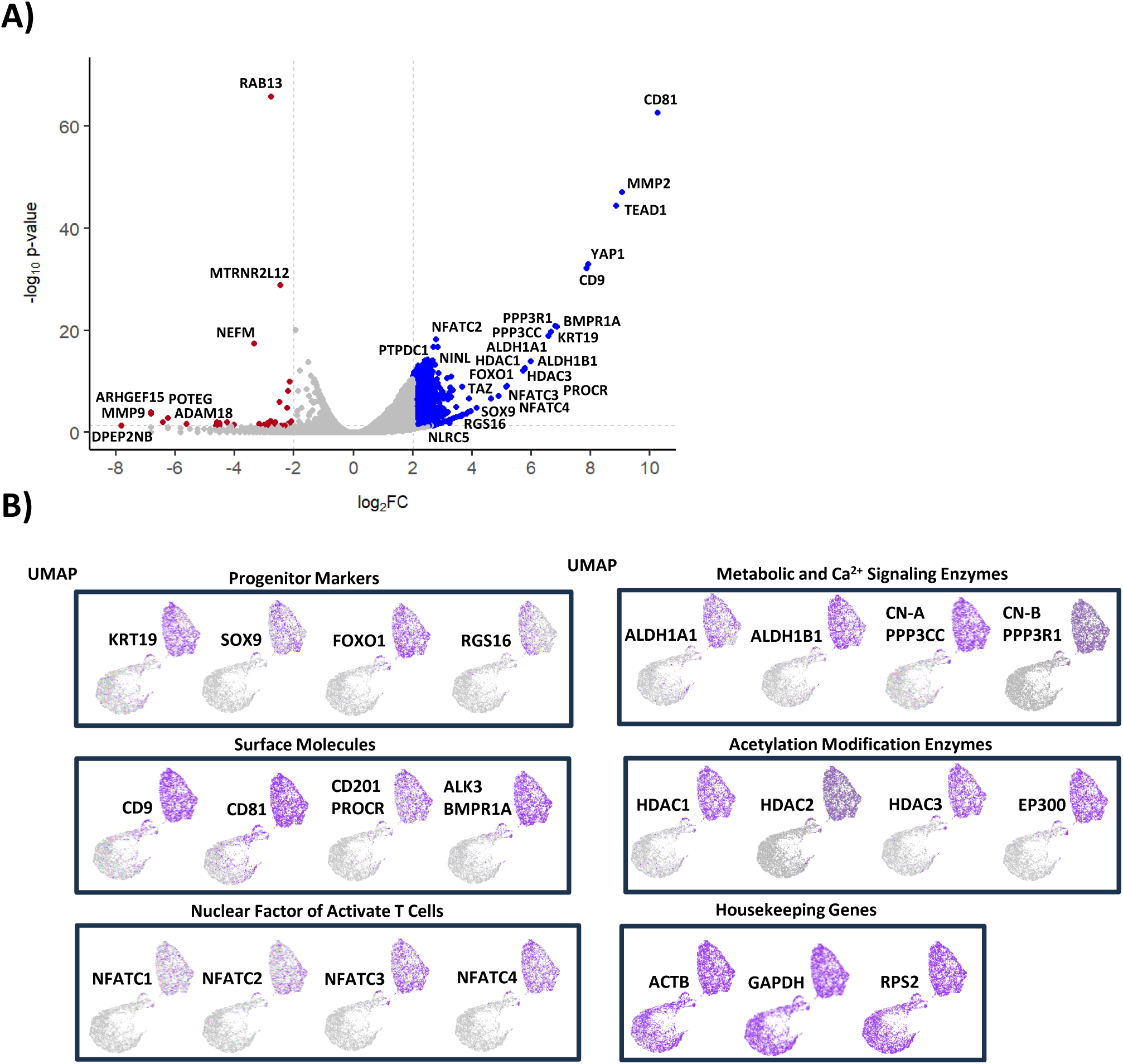
Gene enrichment analysis of IPCs in expanded human pancreatic tissue. (A) Differential gene expression analysis and (B) Uniform Manifold Approximation and Projection (UMAP) analysis of IPCs. Data shown are results from one donor.

Uniform Manifold Approximation and Projection (UMAP) highlighted selective expression of progenitor genes *KRT19*, *SOX9*, *FOXO1*, and *RGS16*, enriched in one of two major IPC subpopulations (Fig. 2B). This cluster also co-expressed surface molecules (*CD9*, *CD81*, *PROCR/CD201*, *BMPR1A*), metabolic regulators (*ALDH1A*, *PPP3CC*), histone modification enzymes (*HDAC1-3*, *EP300*), and family transcription factors (*NFATC1–C4*).

The gene enrichment analyses suggest that the expanded IPC population is in a state of EMT, as previously shown to be a characteristic of an islet cell progenitor state (56–58). Moreover, IPCs selectively express surface markers previously identified in pancreatic endocrine progenitor cells and immature beta cells, including CD9, CD81, PROCR, and BMPR1A (34, 59–63). The IPCs are enriched with ALDH1 isozymes 1A1 and 1B1, which were shown to be associated with regulation of differentiation of endocrine progenitor cells (64, 65). High relative expression of histone modification enzymes p300 and HDAC family members indicates the cells’ potential capability to regulate genes at the chromatin level. The IPCs are also enriched with calcium/calmodulin dependent phosphatase calcineurin (CN) A and B subunits and downstream transcription factor targets of the NFAT family, suggesting potential capacity to modulate genes in response to calcium signaling. This is in line with previous observations that beta cells utilize CN/NFAT signaling to maintain beta-cell differentiation, mass, and function (47, 66–72). Altogether, these data indicate that the expanded IPCs express gene profiles with islet endocrine progenitor characteristics and the potential capacity to support islet cell regeneration and function.

### IPCs Co-transplanted with Islets Enhance Graft Function and Partially Restore Native Pancreatic Endocrine Function in Diabetic Nude Mice

Several studies have shown that ductal epithelial cells derived from pancreatic tissue contain cells that can differentiate into insulin-producing cells (24, 73). Similarly, cultured mesenchymal stromal cells (MSCs) have demonstrated potential to express insulin in vitro and enhance islet graft outcomes when co-transplanted. Because expanded IPCs exhibit features of both MSCs and ductal epithelial cells, including EMT-associated gene expression, we hypothesized that IPCs might retain islet cell multipotency or confer regenerative support that enhances islet graft function in vivo.

To test this, we co-transplanted expanded human IPCs with a subtherapeutic (marginal) dose of human islets (1500 IEQs) into streptozotocin (STZ)-induced diabetic nude mice. Controls included IPCs alone, marginal islets alone, and a positive control group receiving a full therapeutic dose of 3000 IEQs. Neither IPCs nor marginal islets alone were sufficient to reverse diabetes. However, co-transplantation of IPCs with marginal islets reversed hyperglycemia (<200 mg/dL) for up to 30 days (Fig. 3A). Although one mouse in the co-transplant group showed transient hyperglycemia after 2 weeks, the group average remained comparable to the full-dose islet control (p > 0.05). Intraperitoneal glucose tolerance testing (IPGTT) 30 days after transplantation showed that co-transplanted animals could restore blood glucose to a nondiabetic state within 2 hours of glucose injection in contrast to mice treated with marginal islet doses and IPCs alone (Fig 3B). Upon removal of islet grafts by nephrectomy, mice transplanted with IPCs, marginal dose islets, and full-dose islets alone all lost complete islet cell endocrine function within 3 days (Fig. 3A). However, mice co-transplanted with marginal islets and IPCs maintained significantly lower blood glucose than the other groups. Moreover, co-transplanted mice exhibited impaired glucose tolerance but remained resistant to hyperglycemia and maintained IPGTT profiles comparable to islet control groups (Fig. 3C). These data suggested that the observed residual control of blood glucose in mice co-transplanted with marginal islets and IPCs was attributed in part to restoration of native endocrine function.

**FIGURE 3.**
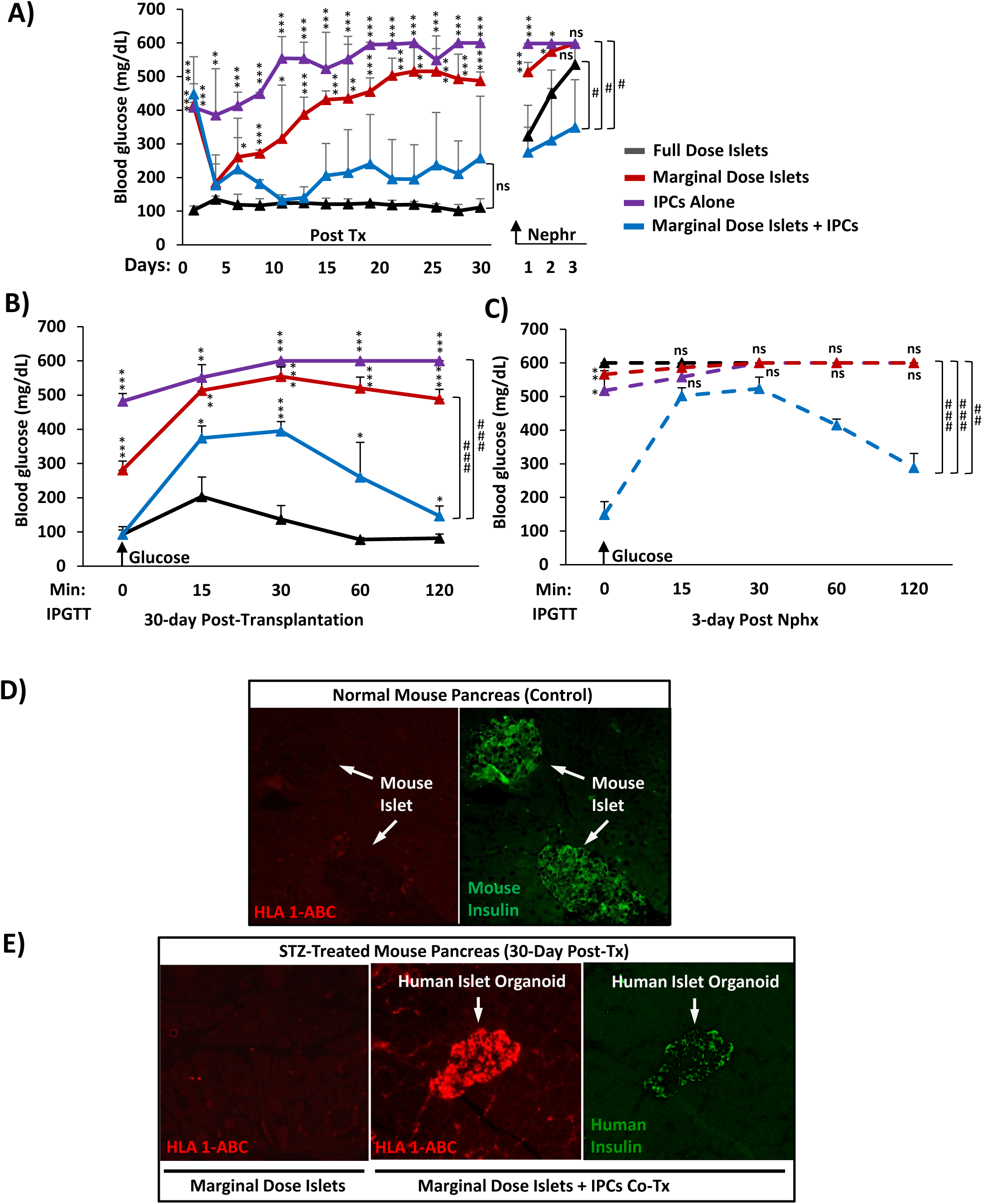
Effects of human IPCs on islet cell function in an STZ-diabetic nude mouse bioassay. Blood glucose profiles of 3000 IEQ full-dose islets (black), 1500 IEQ marginal dose islets (red), IPCs alone (purple), and marginal dose islets co-transplanted with IPCs (blue) for (A) up to 30 days posttransplantation and 3 days postnephrectomy (graft removal) and intraperitoneal glucose tolerance testing (IPGTT) (B) 30 days posttransplantation and (C) 3 days postnephrectomy. Dashed lines represent glucose measurements after islet graft removal. Graphed values are expressed as mean ± SD. Immunofluorescent staining of HLA class 1 molecules A, B, and C (red) and mouse insulin (green) in pancreatic sections procured 30 days posttransplantation from (D) normal mouse pancreas control and (E) STZ-treated diabetic mice transplanted with marginal dose islets alone or with IPCs. Data shown are representative results from at least three independent experiments with 3-5 mice per treatment group.

Because residual control upon graft removal occurred only in mice with islets co-transplanted with IPCs, we reasoned that partial restoration of native endocrine function may have been a result of IPCs contributing to 1) damage repair or regeneration of endogenous mouse islets or 2) direct differentiation of IPCs into functional islet organoids. We therefore examined pancreatic sections of STZ-diabetic mice co-transplanted with marginal islets and IPCs for potential migration of cells by staining for human leukocyte antigen (HLA) class I molecules HLA-A, HLA-B, and HLA-C surface proteins, which are expressed only in human tissue. Diabetic mice co-transplanted with IPCs showed HLA-1 cells scattered throughout the pancreas that were not detected in normal mouse controls (Fig. 3D) or in diabetic mice with human islets transplanted alone (Fig. 3E). Indeed, multiple imaged sections indicated infiltration of human HLA-1^+^ cells that formed islet organoids within the mouse pancreas (Fig. 3B). Notably, insulin production was restricted to HLA-1^+^ islet organoids, indicative of regeneration of human beta-like cells in the mouse pancreas. Human islet organoids were not observed in the pancreas of mice transplanted with islets alone, suggesting that the organoids originated from cells migrating from the human IPC compartment of transplanted cells and did not directly arise from transplanted human islets that may have possibly been dislodged or displaced from the kidney capsule. Collectively, these data demonstrate that IPCs have the capacity to enhance effects of islet cell transplantation to reverse hyperglycemia by improving potency of the islet graft and by restoring native pancreas endocrine function.

### CD9^+^, PROCR^+^ IPCs Form RGS16^+^ IPC Clusters with the Capacity to Differentiate into Islet Organoids in Vitro

Previous studies have indicated that various sources of adult human tissues can differentiate into insulin-producing cells in vitro (25, 28, 31, 43, 44, 74–76). Thus, we sought to determine if expanded IPCs have the potential to differentiate and become glucose-responsive islet-like cells in vitro. Dissociated IPCs formed monolayers that expanded into cellular networks and formed dense islet-like cell clusters upon reaching clonal density within 30 days of culture (Fig. 4A). To define markers for further characterization of IPCs with potential islet regenerative properties, we performed flow cytometry to confirm expression of surface molecules identified in our transcriptomic analysis of highly differentially expressed genes clustering with islet progenitor, immature beta cell, and beta-cell disallowed genes (Fig. 1, 2A). Flow cytometry analysis confirmed co-expression of CD9 and PROCR surface proteins on expanded IPCs (Fig. 4B). The IPCs also internally expressed islet cell progenitor protein RGS16 corresponding to the single cell RNA-seq data, indicating its clustering with islet cell progenitor genes. RGS16 was previously identified as a marker for early pancreatic progenitors that is temporally expressed in IPCs prior to terminal maturation (77, 78). Dimensional reduction analysis heat mapping of protein expression by t-SNE confirmed high co-expression of CD9 and PROCR surface molecules overlapping with a large subset of IPCs expressing RGS16 (Fig. 4C). Immunofluorescent staining showed RGS16 was selectively expressed in the IPC clusters of expanded IPCs which were devoid of insulin (Fig. 5A). Overall, the results suggested that CD9^+^PROCR^+^ IPCs expand and form IPC clusters containing RGS16^+^ islet-like cells in a dedifferentiated or progenitor state.

**FIGURE 4.**
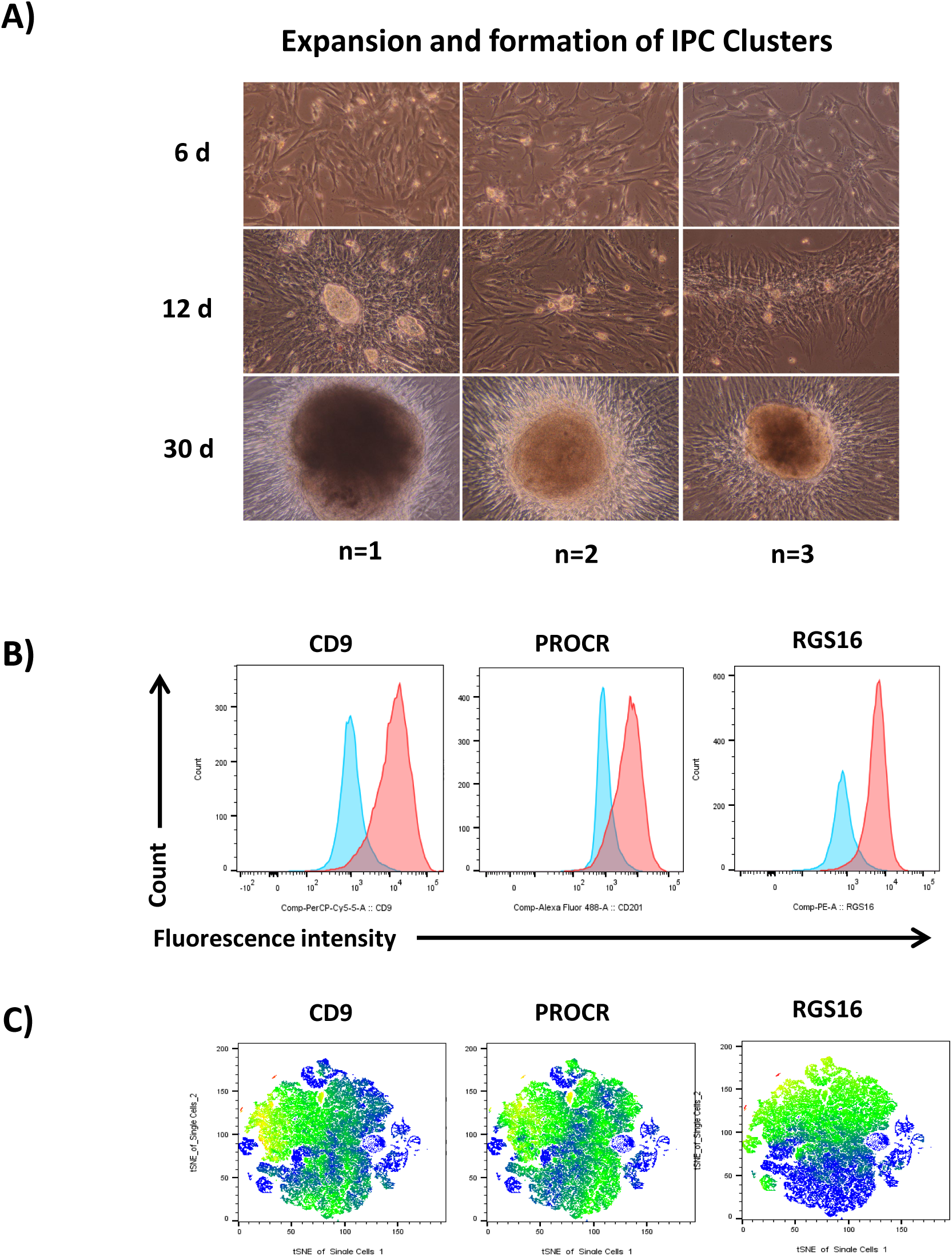
Characterization of IPCs and formation of IPC clusters. (A) Micrograph brightfield (200×) images of expansion of IPCs and formation of IPC clusters within 30 days of culture. Flow cytometry analyses of pancreatic progenitor and beta cell dedifferentiation surface markers CD9 and PROCR and internal islet endocrine lineage marker RGS16 expressed in IPCs by (B) counts and fluorescent intensity and (C) t-distributed stochastic neighbor embedding (t-SNE) heat mapping of CD9, PROCR, and RGS16 protein expression. Data shown are representative results from at least three independent experiments.

**FIGURE 5.**
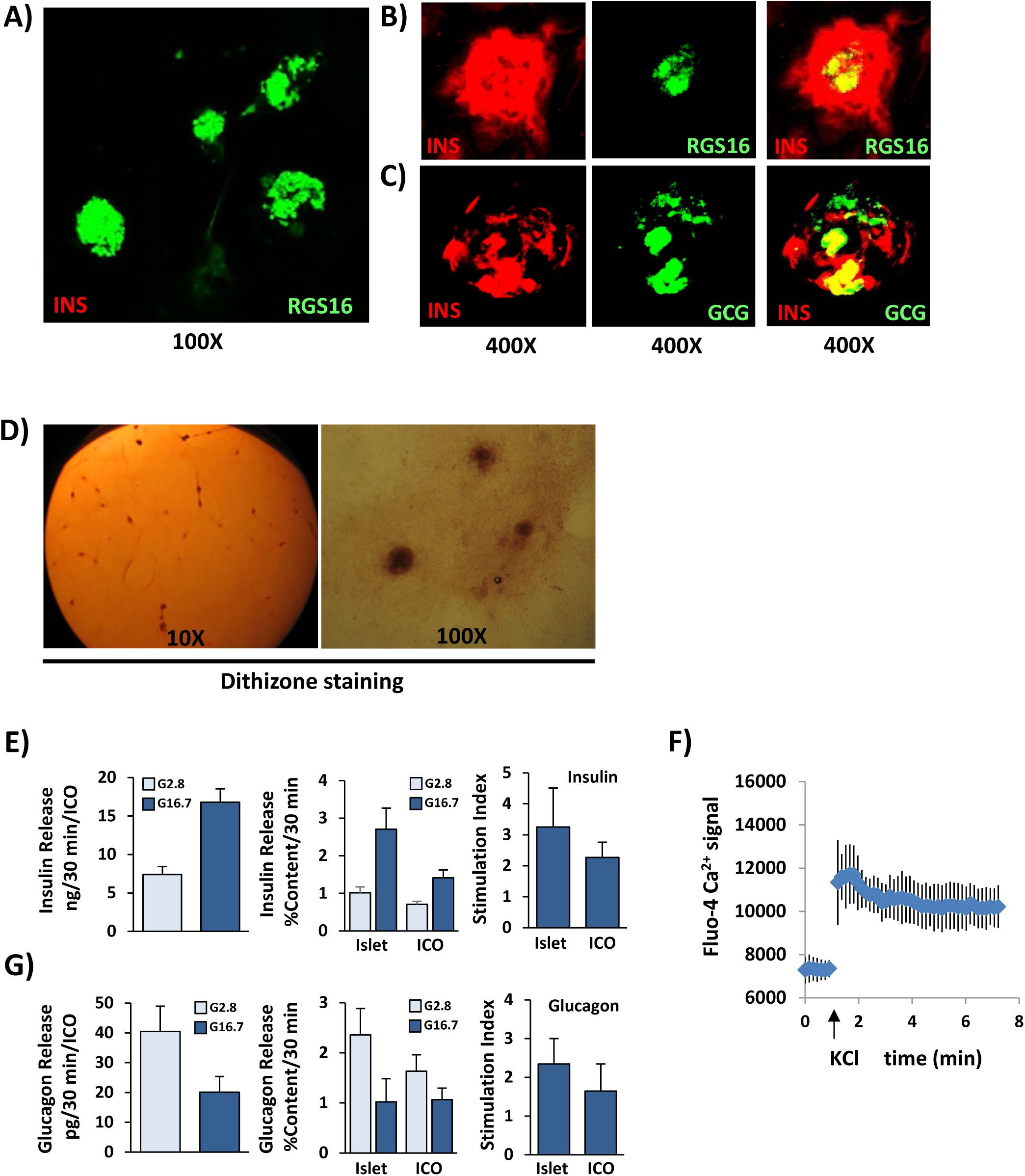
Differentiation of RGS16^+^ IPC clusters into islet organoids by ISX9. Immunofluorescent staining of RGS16 and islet hormones INS and GCG in human IPCs treated with (A) DMSO (control) and (B and C) ISX9 for 14 days in vitro to assess for (B) INS (red)/RGS16 (green) and (C) INS (red)/GCG (green) cellular co-localization. (D) Dithizone staining of IPC organoids treated with ISX9 for 14 days. (E) Glucose-stimulated insulin secretion assay comparing human islet organoids to normal isolated islets. (F) Assessment of intracellular calcium release in islet organoids by KCl-induced depolarization. (G) Glucagon release in islet organoids in response to high (25 mM) and low (2.5 mM) glucose. Graphed values are expressed as mean ± SD. Asterisks above bars indicate statistically significant differences (*p < 0.05, **p < 0.01, ***p < 0.001) in mean values for treatments based on a two-tailed Student’s *t* test. Data shown are representative results from at least three independent experiments.

Our transcriptomic analysis indicated that CD9, PROCR, and RGS16 genes also clustered with cells expressing NFAT and HDAC signaling enzymes. We previously showed that small molecule differentiator isoxazole-9 (ISX9) could restore beta-cell function and prevent dedifferentiation by NFAT-mediated regulation of p300 and HDAC of beta-cell differentiation and disallowed genes at the chromatin level (47). Thus, we hypothesized that ISX9 could likewise induce differentiation or maturation of expanded IPCs that retained this signaling system. Treatment of expanded IPCs with ISX9 for 14 days resulted in selective expression of insulin and glucagon within islet organoids as determined by immunofluorescent staining (Fig. 5B and 5C). Notably, residual RGS16 expression was observed in the inner core of large islet organoids, suggesting that some of the core cells were not completely differentiated (Fig. 5B). Likewise, some core organoid cells co-expressed both insulin and glucagon, indicating that these islet-like cells remained in an immature state (Fig. 5C). However, most insulin-expressing organoid cells no longer expressed RGS16, and a large portion of the islet organoid cells expressed insulin and glucagon independently. Moreover, the islet organoids also intensely stained burgundy red with diphenylthiocarbazone (dithizone), suggesting enrichment of zinc-containing insulin granules, which are produced by mature beta cells (Fig. 5D).

To assess functional maturity of islet organoids, we tested their ability to release insulin and glucagon in response to glucose. Indeed, the islet organoids released insulin in response to high glucose with stimulation indices comparable to those of freshly isolated human islets (Fig. 5E). They also exhibited K^+^-induced depolarization as determined by calcium influx in response to 30 mM K^+^ (Fig. 5F), suggesting functional K_ATP_ channels as expressed in mature beta cells. Furthermore, the islet organoids showed increased glucagon release in response to low glucose, indicative of alpha-like cell function (Fig. 5G). These data indicate that the RGS16^+^ IPC clusters can be differentiated into functional glucose-responsive islet organoids within 14-day treatment with ISX9 in vitro.

### ISX9 Induces a Cascade of Islet Cell Lineage Transcription Factors upon Differentiation to Insulin and Glucagon Expressing Islet Organoids

To identify transcription factors that may contribute to islet organoid differentiation from IPCs in culture, we tracked expression of genes known to be determinants of islet endocrine lineages during defined stages of islet cell development. Time-course analysis showed NFATC2 to be one of the earliest genes expressed, within 6 hours of ISX9 stimulation (Fig. 6). Early endocrine progenitor gene NGN3 was reduced and often undetectable within 14 days of ISX9 treatment. In contrast, islet differentiation transcription factors RFX6 and ND1 were highly inducible within 24 hours, followed by induction of NKX2.2 and NKX6.1 occurring within 48 hours. Beta-cell and alpha-cell restricted transcription factors MAFA and MAFB were expressed within 2 and 7 days, respectively. Finally, insulin and glucagon along with the SLC2A2 (GLUT2) transporter gene expression were detected within 7-14 days upon ISX9 treatment. These data indicate that IPC islet organoid differentiation in vitro recapitulates gene programming similar to what is observed during islet endocrine cell specification in stem cells and developing islets.

**FIGURE 6.**
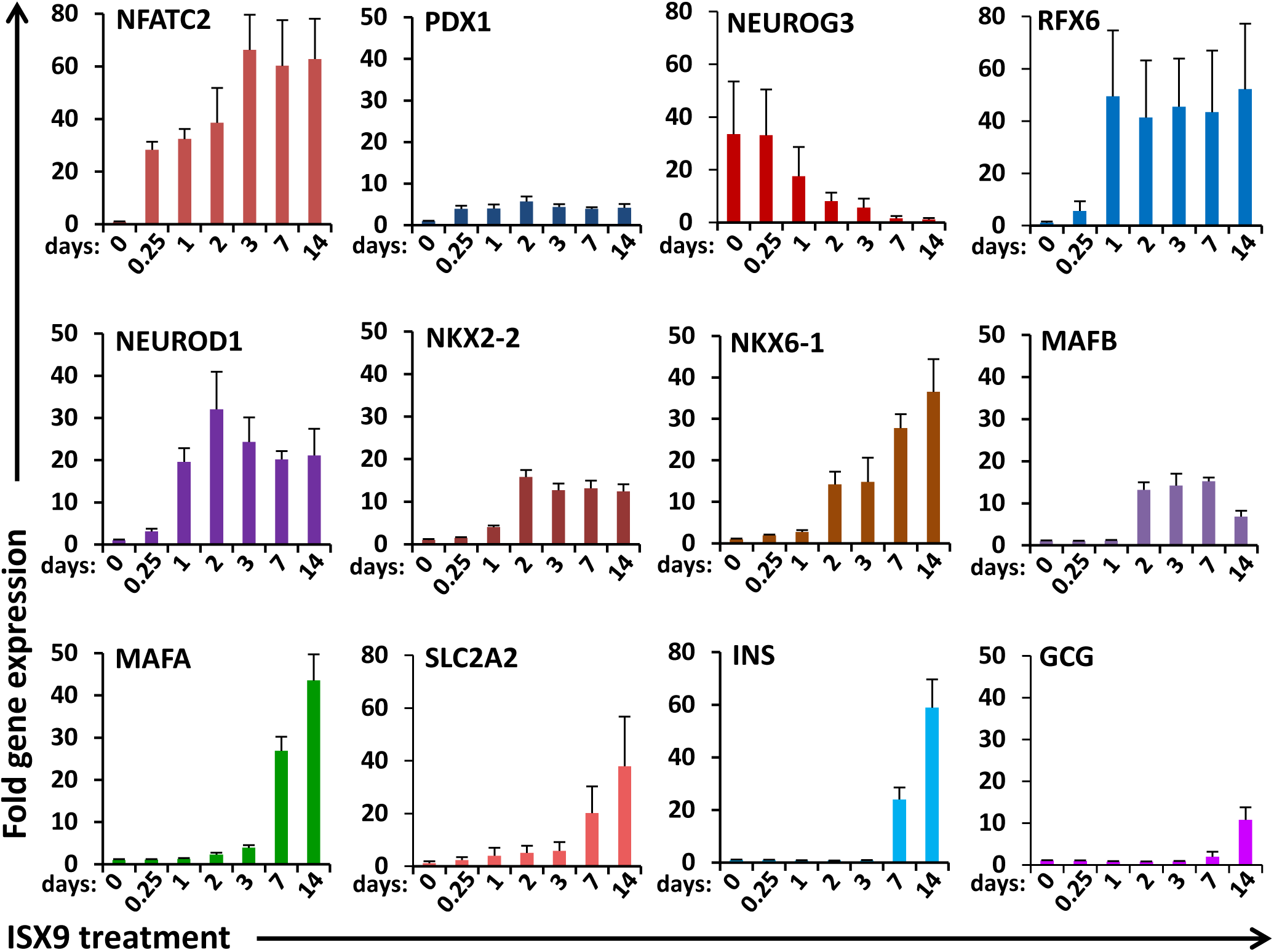
Time-course analysis of islet cell lineage genes expressed during IPC islet organoid differentiation. Expression of mRNA for transcription factors involved in islet cell differentiation and maturation in IPCs treated with ISX9 for up to 14 days. Graphed values are expressed as mean ± SD. Data shown are results from at least three independent experiments.

### CN and NFAT Signaling is Required to Induce RFX6 and NEUROD1 Gene Expression in Differentiating IPC Islet Organoids

RFX6 and NEUROD1 have been shown to be the earliest transcription factors that promote differentiation of endocrine progenitors into islet-specific cell lineages. Both transcription factors are also required to maintain beta-cell identity, maturation, and functional state (79–82). Because ISX9 was previously shown to modulate calcium and CN/NFAT signaling in islet cells, we sought to determine if NFAT was required for activation of RFX6 or NEUROD1 genes during organoid differentiation. ISX9-induced differentiation of IPC organoids correlated with the rapid induction of NFATC2 prior to expression of RFX6 and NEUROD1 in comparison to other NFAT isoforms (Fig. 6, 7A). Selective induction of NFATC2 expression by ISX9 was independent of CN activity, as it was insensitive to CN inhibitor FK506. Promoter-reporter assays showed increased RFX6 and NEUROD1 gene promoter activity after 24-hour treatment with ISX9 (Fig. 7B). Induction of RFX6 and NEUROD1 promoters was blocked by CN inhibitor FK506 or overexpression of a dominant negative NFAT (dnNFAT) protein containing a truncated PxIxIT box motif as compared to DMSO and mutated dnNFAT controls. These data indicate that ISX9 induced expression of both RFX6 and NEUROD1 at the gene promoter level in IPC organoids in a CN-and NFAT-dependent manner.

**FIGURE 7.**
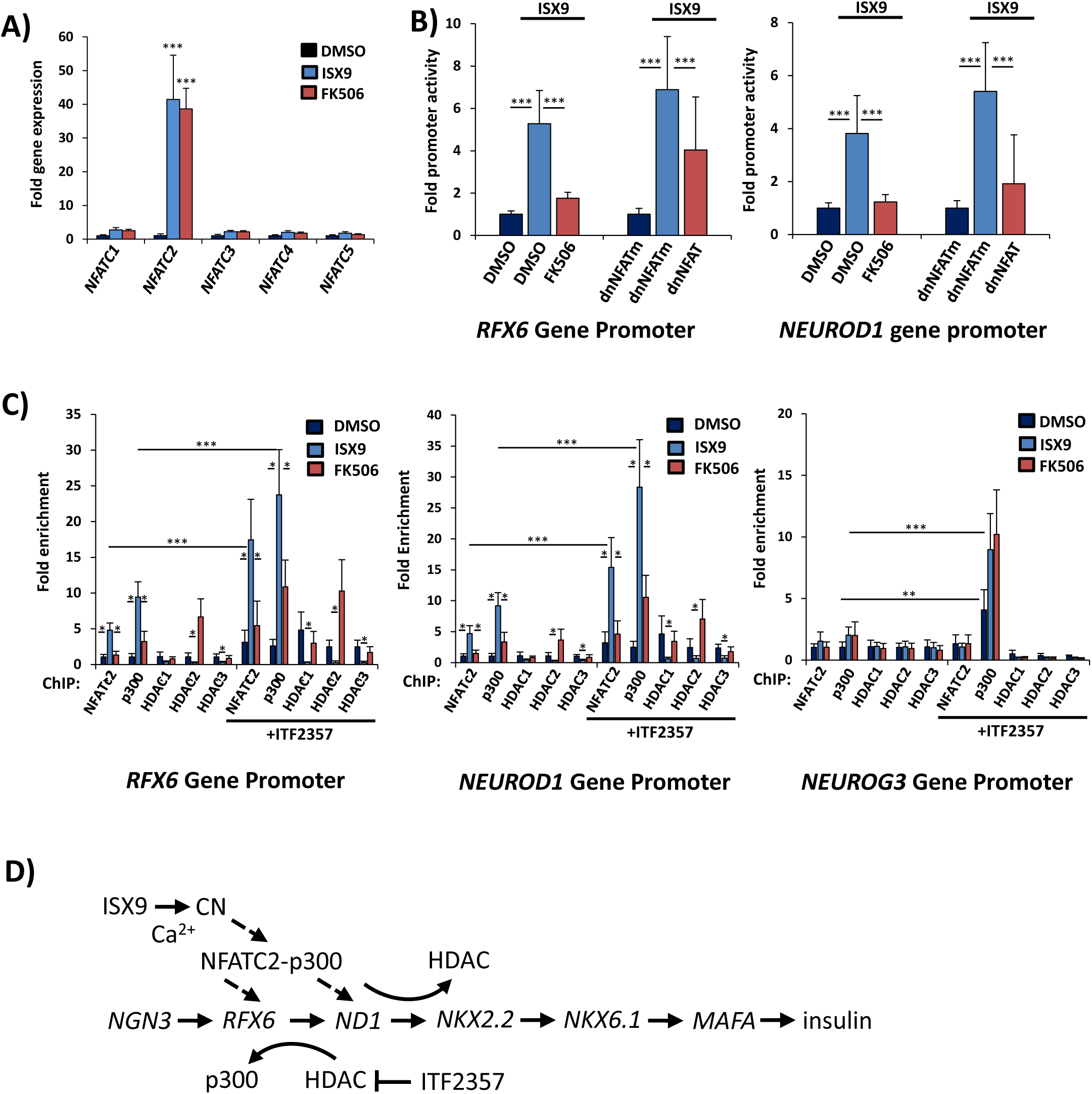
Requirement of CN/NFAT signaling to recruit p300 to RFX6 and NEUROD1 promoters during ISX9-induced IPC islet organoid differentiation. (A) Induction of NFAT family isoform genes in IPC clusters treated with ISX9 for 24 hours in the presence of CN inhibitor FK506. (B) RFX6 and NEUROD1 promoter activation by ISX9 in IPCs in the presence of FK506 or transfected with gene vectors overexpressing dominant negative NFAT (dnNFAT) and mutated control (dnNFATm). (C) ChIP assay of association of NFATC2, p300, HDAC1, HDAC2, and HDAC3 with RFX6, NEUROD1, and NEUROG3 promoters upon 6-hour treatment of IPCs with ISX9 with and without 24-hour pretreatment with ITF2357. (D) Schematic of mechanism of CN/NFATC2 induction of RFX6 and NEUROD1 genes by ISX9 to stimulate islet organoid differentiation. Graphed values are expressed as mean ± SD. Asterisks above bars indicate statistically significant differences (*p < 0.05, ***p < 0.001) in mean values for treatments based on a two-way ANOVA and Sidak’s multiple comparison test. Data shown are results from at least three independent experiments using MSCs derived from three individual donors.

### HDAC Inhibition and Activation of CN/NFAT Signaling Induces Insulin Gene Expression in Adult IPCs

We previously showed that calcium-dependent CN/NFAT is required for beta-cell differentiation and insulin gene transcription (47). Small molecule differentiation inducer ISX9 was able to prevent dedifferentiation of stressed islet beta cells by preserving intracellular calcium signaling and CN/NFATc2-mediated induction of beta-cell transcription factors and repression of disallowed genes. In contrast, extended exposure of islets to stress resulted in loss of CN/NFATc2 signaling, accumulation of HDACs on the RFX6 gene promoter, and beta-cell dedifferentiation. Thus, we hypothesized that inhibiting HDACs and reactivating CN/NFAT in IPC clusters would promote their differentiation into islet organoids.

To test this hypothesis, we pretreated IPCs with HDAC inhibitor ITF2357 for 24 hours prior to treatment with ISX9 or inhibition of NFAT deactivating kinase Dyrk1a by harmine. Insulin gene transcription was induced within 10 days of exposure to both ISX9 and harmine, which in each case was accentuated by pretreatment with ITF2357 (Suppl. Fig. 1). Both ISX9 and harmine showed enhanced effects to induce insulin expression in IPCs compared to treatment with modified end-stage differentiation protocols commonly used to stimulate endocrine cell progenitor stage to mature sc-islets (stages 5-7) in hESCs and iPSCs. These results suggest that CN/NFAT signaling can be targeted to pro-differentiate adult IPCs with relatively high proficiency.

### NFATc2 Mediates Chromatin Assembly of Histone Acetylation-Modifying Enzymes p300 and HDACs on the RFX6 and NEUROD1 Gene Promoters

To determine if NFATC2 directly binds to the RFX6 gene promoter, we performed chromatin immunoprecipitation (ChIP) assays on islet organoids cultured in ISX9 for 24 hours. ChIP analysis revealed significant increases in enrichment of NFATC2 on the 5ʹ flanking region of the RFX6 and NEUROD1 gene promoters (Fig. 7C). In contrast, ISX9 had no effect on NFATC2 binding to the upstream endocrine progenitor transcription factor NEUROG3. Association of NFAT with the RFX6 and NEUROD1 promoters was largely prevented by FK506, indicating the requirement of CN for NFAT-mediated transcription. Previous studies showed that ISX9 could enhance p300 acetyltransferase recruitment to the insulin gene promoter to acetylate histones in MIN6 beta cells [41]. We therefore sought to determine effects of ISX9 to target chromatin-related proteins to the RFX6 and NEUROD1 gene promoters in islet organoids. ChIP analysis was performed on ISX9-treated islet organoids to assess DNA promoter assembly of NFATC2 and histone-modifying enzymes p300 and HDACs. NFATC2 association correlated with accumulation of p300 upon the RFX6 and NEUROD1 promoters within 6 hours of ISX9 treatment (Fig. 7C). By contrast, HDACs were depleted on the RFX6 and NEUROD1 promoters under conditions that stimulated NFATC2 binding. In each case, FK506 inhibited effects of ISX9 to promote p300 and diminish HDAC accumulation on the RFX6 promoter. In contrast, ISX9 and FK506 did not affect association of p300 or HDACs with the NGN3 promoter, indicating that ISX9-induced CN/NFAT signaling mediates specificity of p300 recruitment toward the RFX6 and NEUROD1 genes (Fig. 5D). Pretreatment of IPC clusters with HDAC inhibitor ITF2357 (50 nM) enhanced the effects of ISX9 to recruit p300 to the RFX6 and NEUROD1 promoters and also induced a significant increase in the ratio of p300:HDAC accumulation on the NEUROG3 promoter. FK506 blocked effects of ITF2357 on the RFX6 and NEUROD1 promoters, but p300 and HDAC accumulation on the NGN3 promoter were unchanged. Increases in p300:HDAC ratios correlated with increased gene expression. Whereas ISX9 selectively induced transcriptional activity of the RFX6 and NEUROD1 genes, ITF2357 globally promoted transcription from all genes. The results delineate CN/NFAT-mediated displacement of HDACs by p300 to selectively activate the RFX6 and NEUROD1 genes in response to ISX9 and a global effect of ITF2357 to increase the ratio of p300:HDAC promoter accumulation on genes to enhance induction of transcriptional activity. These data suggest that CN/NFATc2-mediated recruitment of p300 and induction of the RFX6 and NEUROD1 genes bypasses requirements of NGN3 to promote differentiation of IPCs to an islet-like cell type (Fig. 7D).

### IPC-derived Islet Organoids are Glucose- and GLP-1-Responsive

To release appropriate and sustainable insulin in response to physiological demand, beta cells must regulate insulin production in response to changes in glucose and nutrient-responsive incretins including peptide hormone glucagon-like peptide 1 (GLP-1). Therefore, we assessed effects of high (16.7 mM) glucose and GLP-1 on insulin gene expression and output in IPC-derived islet organoids (Fig. 8). ChIP analysis of the insulin promoter demonstrated binding of key transcription factors PDX1, NEUROD1, and MAFA known to regulate insulin gene transcription in mature beta cells (Fig. 8A). NFATC2 and MAFA showed highest enrichment upon the insulin gene promoter in response to glucose and GLP-1. The effect of NFATC2 and MAFA was enhanced along with significant enrichment of PDX1 and NEUROD1 upon the insulin gene promoter when IPC clusters were primed for 24 hours with ITF2357 prior to 24-hour treatment with ISX9. Islet organoids transfected with a luciferase insulin gene promoter-reporter showed upregulation of the insulin gene in response to glucose and GLP-1, corresponding to increased binding of NFATc2 with PDX-1, NeuroD1, and MafA to the insulin gene promoter (Fig. 8A, 8B). Insulin promoter activity was stimulated more than 5-fold in the presence of high glucose and GLP-1, which was enhanced to more than 10-fold when IPC clusters were primed with ITF2357. To test effects of glucose and GLP-1 on insulin secretion, we performed glucose-stimulated insulin secretion assays on IPC-derived islet organoids. High glucose treatment of islet organoids for 2 hours resulted in insulin release of 7.7 ± 2.1 pg/h/organoid and 17.0 ± 5.3 pg/h/organoid in the presence of GLP-1 (Fig. 8C). Overall, these data indicate that differentiated islet organoids produce and release insulin in response to glucose and GLP-1 by mechanisms similar to those observed in mature pancreatic beta cells.

**FIGURE 8.**
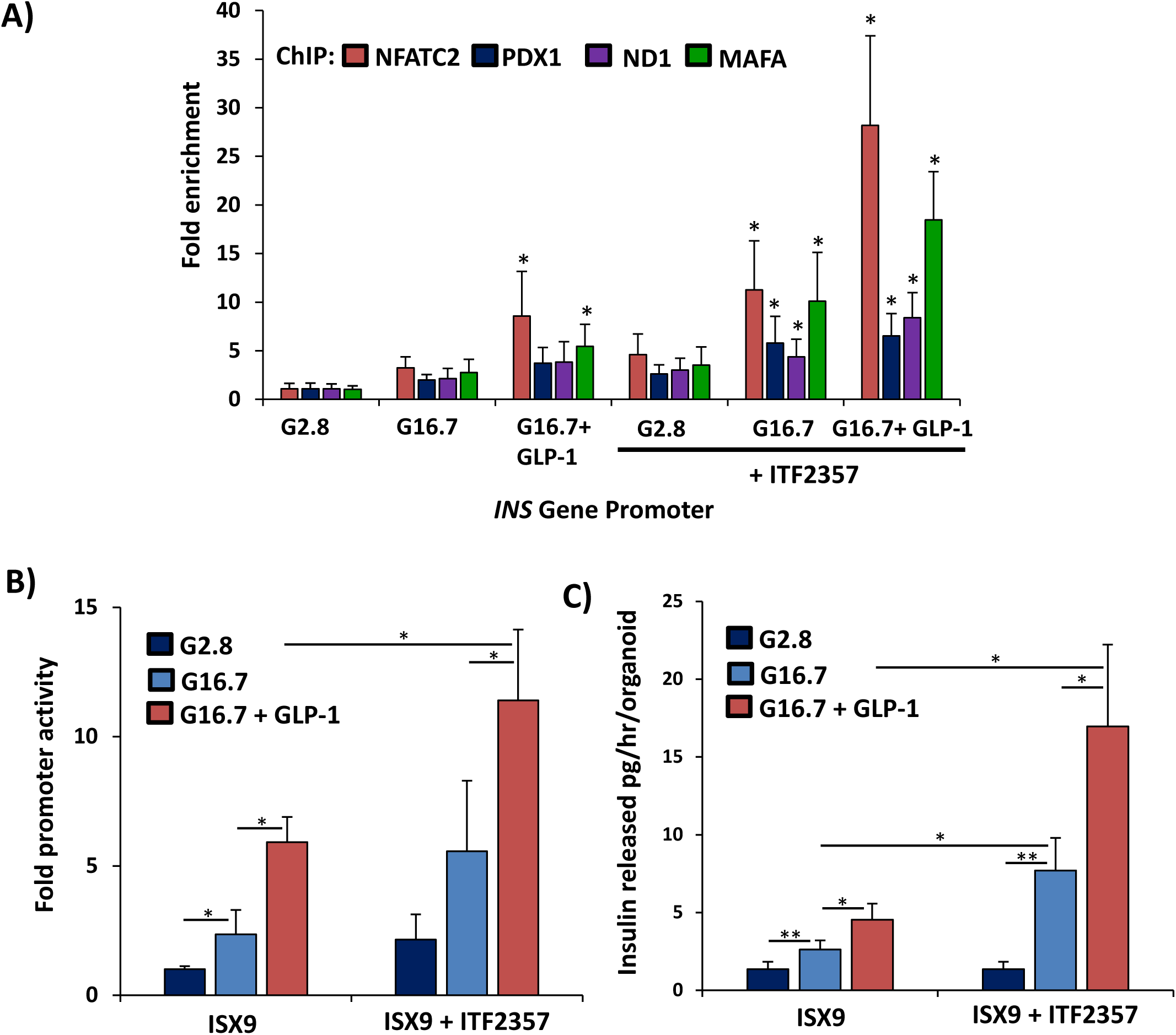
Glucose and GLP-1 responsiveness of differentiated IPC islet organoids. (A) ChIP assay of association of NFATC2, PDX1, ND1, and MAFA in islet organoids in response to 20-minute treatment with high glucose (G16.7) and GLP-1. (B) Insulin-promoter activity in response to 24-hour treatment with high glucose (G16.7) and GLP-1. (C) Glucose-stimulated insulin secretion assay of islet organoid insulin secretion in response to 2-hour treatment with high glucose (G16.7) and GLP-1. Graphed values are expressed as mean ± SD. Asterisks above graphs indicate statistically significant differences (*p < 0.05) in mean values for treatments compared to 2.8 mM glucose controls based on a two-tailed Student’s *t* test. Asterisks above bars indicate statistically significant differences (*p < 0.05, **p < 0.01) in mean values for treatments based on a two-way ANOVA and Sidak’s multiple comparison test. Data shown are results from at least three independent experiments using islet cultures derived from three individual donors.

## DISCUSSION

A large body of evidence over the past century has indicated that the pancreas possesses regenerative capacity for both exocrine and endocrine tissue (83–89). Several models of pancreatic injury have shown that repair and restoration are facilitated in part by stem cell progenitors, which are capable of regenerating pancreatic tissue (83–90). However, the identity of adult islet progenitor cells remains unresolved, primarily due to the lack of specific markers for defining or tracking such populations. Moreover, the notion that adult stem/progenitor cells contribute to islet regeneration has been controversial, as lineage tracing studies in transgenic mice have suggested that increases in beta-cell mass after birth occur primarily through replication of pre-existing beta cells (40).

In this study, we identified and characterized a subset of expandable adult pancreatic cells—CD9⁺, PROCR⁺, RGS16⁺ islet pancridia cells (IPCs)—that give rise to IPC clusters with the capacity to differentiate into glucose-responsive islet organoids in vitro and in vivo. Using this model, we further explored the mechanisms governing islet lineage differentiation by interrogating developmental transcriptional programs activated during IPC-to-organoid transition.

Differentiation of IPC clusters into insulin- and glucagon-producing islet organoids could be driven by ISX9, a small molecule previously shown to preserve beta-cell identity through chromatin modulation. ISX9-induced differentiation was dependent on calcineurin (CN) signaling and binding of NFATC2 to the RFX6 and NEUROD1 gene promoters. This promoter association was accompanied by CN-dependent recruitment of the histone acetyltransferase p300 and displacement of histone deacetylases HDAC1–3, suggesting that transcriptional activation is regulated, at least in part, by site-specific chromatin acetylation. Pretreatment with the HDAC inhibitor ITF2357 globally increased p300 enrichment on the *RFX6* promoter and enhanced islet organoid differentiation.

Collectively, these data suggest that CN/NFATC2 contributes to selective epigenetic induction of the RFX6 gene to bypass the need for *NGN3*, a transcription factor essential during embryonic islet development [6, 7]. In contrast, induction of the NGN3 promoter to enhance the potency of differentiation by HDAC inhibition occurred independently of CN activity or binding of NFATC2. These distinct mechanisms define critical signaling components required for inducing genes that initiate differentiation of IPC clusters into islet organoids.

Importantly, induction of the RFX6 and NEUROD1 genes permitted expression of genes in islet organoids required for both terminally differentiated beta cells and alpha cells. The resulting mature islet organoids expressed and released insulin and glucagon in a glucose-regulated manner. The islet organoid beta-like activated canonical insulin transcription factors, including PDX1, NEUROD1, and MAFA, and exhibited NFATC2 binding to the insulin promoter in response to glucose and GLP-1, recapitulating signaling pathways used by mature beta cells to maintain glucose responsiveness.

Overall, these findings provide new insight to methodologies and mechanisms for expansion and differentiation of adult human IPCs into islet organoids. While recent advances have demonstrated the ability to derive beta cells from human pluripotent stem cells (1, 2, 91–93), no naturally existing adult islet progenitor cell has been definitively identified to selectively regenerate islet organoids in humans. We report that IPC clusters selectively express RGS16, an islet progenitor marker, and can be differentiated by ISX9 via CN/NFAT-mediated activation of *RFX6*. This process is enhanced by ITF2357, which facilitates chromatin remodeling by NFAT to induce activation of genes required for driving islet cell differentiation and maturation.

The identification of signaling pathways and transcriptional targets that govern adult IPC differentiation into functional islet organoids represents a critical advance toward regenerative medicine strategies for diabetes. These findings may enable the development of scalable, cell-based and pharmacologically directed therapies to restore islet endocrine function in individuals with pancreatic disease or diabetes.

## MATERIALS AND METHODS

### Reagents and recombinant DNA constructs

Antibodies used were as follows: NFATC2, RGS16, NEUROD1, NEUROG3, p300, HDAC1, HDAC2, and HDAC3 (Santa Cruz Biotechnology, Inc); CD45-Alexa Fluor and CD29-Alexa Fluor (BioLegend); HLA Class 1 ABC (Abcam); and CD34-APC, CD105-APC, CD90-PE, CD73-PE (BD Biosciences). Gluc-ON reporters for RFX6 (HPRM53326-PG04), NEUROD1 (HPRM69533-PG04), and INS (HPRM30189-PG04) promoters were obtained from GeneCopoeia. Plasmid expression vectors dominant-negative NFAT PxIxIT motif (dnNFAT) and mutated dnNFAT AxAxAA motif (dnNFATm) were previously described (68, 94–96).

### Cell and tissue culture

Digested pancreatic tissue from multiple donors was obtained from the cGMP Islet Cell Processing Laboratory at Baylor University Medical Center under Institutional Review Board-approved protocols in accordance with institutional and national guidelines and regulations. Pancreatic tissue and IPCs were washed, cultured, and expanded in RPMI 1640 (Gibco) containing 11 mM glucose, 10% heat-inactivated fetal bovine serum, 10 mM HEPES (pH 7.4), 2 mM L-glutamine, 1 mM sodium pyruvate, 50 μM β-mercaptoethanol, 100 U/mL penicillin, and 100 μg/mL streptomycin at 37°C in 5% CO_2_ humidified air. Islet progenitor cells were expanded to high clonal density for up to 28 days to produce IPC clusters. Analysis of IPCs and IPC clusters was performed on weekly passages 8-15. FBS was reduced to 2-5% for ISX9-induced islet cell differentiation. Krebs-Ringer bicarbonate HEPES buffer media was used with low (2.8 mM) and high (16.7 mM) glucose for glucose-stimulated insulin secretion and high (50 mM) KCl for depolarization experiments. RPMI 1640 with 2 mM L-glutamine, 1:200 ITS-X, 10 µM triiodo-L-thyronine (T3), 10 µM ALK5 inhibitor II, 10 µM ZnSO_4_, and 10 µg/mL heparin was used as a base medium for end stages 5-7 of sc-islet differentiation supplemented as follows: stage 5 endocrine progenitors (SANT-1, 0.05 µM retinoic acid, and 100 nM LDN193189 for 3 days), stage 6 immature sc-islets (100 nM gamma secretase inhibitor XX (GSiXX), and 100 nM LDN193189 for 7 days), and stage 7 mature sc-islets (n-acetyl cysteine (NAC), Trolox, and 2 μM R428 for 7d).

### Sc-RNA sequenzcing and analysis

IPCs were digested to single-cell suspensions and libraries were constructed with DNA barcodes and unique molecular index (UMI) adapters using the 10x Genomics Chromium Next GEM Single Cell 3’ Reagent Kit Version 3. Samples were loaded into 10x chip wells to produce Gel Bead-in-Emulsions. Libraries were sequenced using a P2-100 flowcell on an Illumina NextSeq 2000 platform. A total of 100,000 barcodes were filtered and aligned to human genome assembly reference GRCh38-2020-A using the Cell Ranger Count v7.1.0 pipeline with default parameters. Automated reference-based processing and mapping to integrated pancreas reference datasets was performed by Azimuth (HuBMAP). Seurat and Loupe Browser were used for data integration, analysis, and visualization to generate 20 cell clusters. Cells were restricted to 200-82150 UMIs with at least 200-8749 detected genes and less than 20% expressed mitochondrial genes. Cell clusters enriched with islet progenitor gene markers were further selected for analysis.

### DNA transfection

Intact islet organoids were detached from tissue culture by 10-minute incubation with TrypLE Express (Gibco) at 37°C and filtered prior to electroporation of ∼100 islet organoids per sample with 2 µg total DNA using a Neon Transfection System (ThermoFisher Scientific) for 2 pulses of 1200 V for 20 ms. Islet organoids were cultured 24 hours prior to experimental treatments. The Secrete-Pair Dual Luminescence Assay Kit (GeneCopoeia) was used to normalize for promoter-reporter transfection. Cells were then lysed in passive lysis buffer, and Firefly and Renilla luminescence was measured with a dual luciferase assay kit (Promega). Luminescence was measured using the Cytation5 Cell Imaging Multi-Mode Reader (BioTek).

### Diabetic nude mouse bioassay

Nude mice were rendered diabetic by intraperitoneal administration of 200 mg/kg body weight of STZ (Sigma). Mice were considered diabetic when fasting blood glucose was >200 mg/dL for 3 or more consecutive days. Full-dose human islets (3000 IEQ), marginal-dose human islets (1500 IEQ), IPCs alone (1×10^6^), and marginal-dose islets with IPCs were transplanted into kidney capsules of diabetic nude mice. Blood glucose and animal health were monitored daily. Mice were fasted for 6 hours with water provided ad libitum prior to intraperitoneal administration of 20% glucose solution at 2 g glucose/kg body weight. Blood glucose was measured at 30-minute intervals for intraperitoneal glucose tolerance test (IPGTT) analyses. All animal surgeries and procedures were performed in accordance with approved institutional animal care and use committee protocols.

### Immunofluorescent staining

Paraffin-embedded tissue sections from mouse pancreases and islet-kidney grafts were washed with PBS containing 0.1% Triton X-100 and 4% BSA. Primary antibody (1:250) in the same solution was incubated with cells overnight at 4°C. After washing, cells were incubated with secondary antibody (1:5000) in PBS containing 0.1% Triton X-100 and 1% BSA for 1 hour. Samples were imaged and analyzed by an Olympus BX61 TRF Fluorescent Microscope.

### Flow cytometry

Cells were detached from the culture flask and dissociated into single cells by 10-minute incubation in TrypLE Express (Gibco) at 37°C. Dissociated IPCs were filtered through a 40 µm sterile nylon cell strainer (Fisher Scientific), and single cells were then washed twice with FACS buffer (2% FBS in PBS). Viability and counting were performed using 0.4% Trypan Blue solution (Thermo Fisher Scientific). A total of 1 × 10^6^ cells in suspension were labeled with fluorochrome-conjugated surface antibody CD9. For intracellular staining of PROCR and RGS16, cells were fixed and permeabilized using Cytofix/Cytoperm (BD Biosciences) according to the manufacturer’s instructions. FACS data were acquired on an LSR Fortessa flow cytometer with 5 lasers and 18 channels (BD Biosciences). Fluorescence minus one (FMO) cell sets and unstained cells served as biological negative controls. FACS data were analyzed by FlowJo software version 10.10.0.

### Chromatin immunoprecipitation (ChIP) assays

ChIP assays were performed as previously described (47). Islet organoids were fixed, and chromatin DNA-protein was cross-linked with 1% formaldehyde and sonicated with a Bioruptor 200 (Diagenode) to produce DNA fragments. DNA-protein complexes were immunoprecipitated with indicated antibodies or IgG isotype controls, extensively washed, and the cross-links were reversed by heating to 65°C for 4 hours in Tris-HCl (pH 6.5), 5 M NaCl, and 0.5 M EDTA. DNA was extracted by phenol/CHCl3 and precipitated with ethanol. Precipitated DNA and 1% control inputs were analyzed by real-time quantitative polymerase chain reaction (qPCR).

### Real-time qPCR

Total RNA was isolated with the Qiagen RNeasy Mini Kit. Equal amounts of cDNA were synthesized using a High Capacity Reverse Transcription Kit (ThermoFisher). TaqMan primers were mixed with TaqMan Universal Master Mix II, with uracil N-glycosylase (ThermoFisher). 18S was amplified as an internal control. qPCR was performed using the Bio-Rad CFX connect system with TaqMan Primer Assays to detect 18S and target genes (ThermoFisher). RT² qPCR primer assays were used to amplify NFATC2, PDX1, NEUROG3, RFX6, NEUROD1, NKX2-2, NKX6-1, MAFA, MAFB, SLC2A2, INS, and GCG genes (Qiagen).

Primer sequences used to detect DNA input and immunoprecipitated 5′-flank promoter regions of human NEUROG3, RFX6, NEUROD1, and INS genes included RFX6 -499 to -274, 5ʹ-TGCAAAGACTGAGCGGTACT and 5ʹ-CCCTCCTTCCCGTTCTCTCA; NEUROD1 -516 to - 70, 5ʹ-AGGCCACTCGCTCTGATCTA and 5ʹ-CTGAGGGGCTAGCAGGTCTA; NEUROG3, to -598 to -21, 5ʹ-TGGAAGGGACATAGGCAGGA and 5ʹ-CACGCTTTATCTGCTTCGCC; and INS -204 to -46, 5′-GTCCTGAGGAAGAGGTGCTG and 5′-CCATCTCCCCTACCTGTCAA.

### Insulin and glucagon secretion assays

Islet clusters pretreated with ITF2357 for 24 hours and ISX9 for 14 days were isolated from cell cultures and incubated for 2 hours in DMEM medium for recovery. Islet organoids were then transferred to low (2.8 mM) or high (16.7 mM) glucose in KRBH medium for 60 minutes before replacing and stimulating in static incubation conditions with high (for insulin release) or low (for glucagon release) glucose KRBH medium for 60 minutes, respectively. Insulin and glucagon protein from cells and supernatant were measured by Insulin (ALPCO) and Glucagon Quantikine (R&D Systems) ELISA kits, respectively.

### Statistical analysis

Statistical analysis was performed using GraphPad Prism version 9.0 (La Jolla, CA). Statistical significance for more than two groups was determined by two-way analysis of variance and Sidak’s multiple comparison test. Direct comparisons between two experimental groups were determined using unpaired Student’s *t* test. Differences were considered significant when *p* values were < 0.05 (*), < 0.01 (**), and < 0.001 (***).

## ACKNOWLEDGMENTS

We thank Jean Philippe Blanck, Rauf Shahbazov, Faisal Kunnathodi, and Prathab Balaji Saravanan for technical assistance. We thank Ashley Hopkins at Baylor Scott & White Research Institute for administrative support throughout the development and filing of the related patent application. We also acknowledge the Baylor Healthcare System Foundation for funding support.

