## Supplemental Tables and Figure 1 for "Identification and Characterization of Adult Islet Pancridia Cells Capable of Differentiating into Islet Organoids"

**Suppl Table 1**

| Human pancreas donor characteristics | | | | |
|---|---|---|---|---|
| **Identifier/UNOS#** | Age | Sex | BMI | Ethnicity |
| **Donor_001** | 65 | Female | 24.2 | White |
| **Donor_002** | 57 | Female | 23 | White |
| **Donor_003** | 45 | Male | 29.8 | White |
| **Donor_004** | 25 | Female | 35.5 | Black |
| **Donor_005** | 22 | Male | 33.2 | Hispanic |
| **Donor_006** | 29 | Male | 26.3 | White |
| **Donor_007** | 60 | Male | 26.7 | Black |
| **Donor_008** | 60 | Male | 28.7 | Hispanic |
| **Donor_009** | 46 | Male | 30.7 | Hispanic |

### Suppl Table 2

**Markers used to identify cel types**

| Cell Type | Cell Markers |
|---|---|
| Acinar Cell | AMY2A, AMY2B, CPA1, PTF1A |
| Islet Cell | INS, GCG, IAPP, SST, PPY |
| Islet Progenitor Cell | PROCR, BMPR1A, RGS16, ISL1, NES |
| Immature Beta Cell | CD9, CD81, YAP1, MMP2, FOXO1, HK1 |
| Beta Cell Disallowed | LDHA, SCL16A1, ACOT7, OAT, PDGFRA, CAT, IGFBP4, ZFP36L, ZYX, LMO4 |
| Ductal Epithelial | KRT19, SOX9, CFTR, EPCAM, CDH1 |
| MSC Positive | ENG (CD105), NT5E (CD73), THY1 (CD90), ITGB1 (CD29) |
| MSC Negative | PTPRC (CD45), CD34, CD14, ITGAM (CD11b), CD79A, CD19, HLA-DRA |
| Stellate Cell | ACTA2 (alpha-SMA), CYGB |
| Endothelial Cell | CDH5, PECAM1, VWF, CLDN5 |

### Suppl. Table 3

**Top cell type enrichment genes - PanglaoDB Augmented**

| Index | Name | P-value | Odds Ratio | Combined score |
|---|---|---|---|---|
| 1 | Pancreatic Progenitor Cells | 0.00002316 | 11.70 | 124.85 |
| 2 | Osteocytes | 0.0001328 | 11.21 | 100.06 |
| 3 | Kidney Progenitor Cells | 0.0001837 | 10.42 | 89.64 |
| 4 | Cardiac Stem And Precursor Cells | 0.0002588 | 9.65 | 79.67 |
| 5 | Adipocyte Progenitor Cells | 0.0002807 | 9.47 | 77.44 |
| 6 | Luteal Cells | 0.0004124 | 8.68 | 67.61 |
| 7 | Pulmonary Alveolar Type I Cells | 0.0005111 | 8.26 | 62.60 |
| 8 | Melanocytes | 0.0005477 | 8.13 | 61.05 |
| 9 | His Bundle Cells | 0.001483 | 8.78 | 57.18 |
| 10 | Embryonic Stem Cells | 0.001913 | 6.07 | 38.01 |

**Top cellular and tissue enrichment genes - CellMarker 2024**

| Index | Name | P-value | Odds Ratio | Combined score |
|---|---|---|---|---|
| 1 | Progenitor Cell Pancreas Human | 0.0006796 | 67.67 | 493.57 |
| 2 | Ductal Cell Pancreas Human | 0.00006478 | 47.31 | 456.30 |
| 3 | Plasma Cell Undefined Human | 0.00009371 | 41.00 | 380.29 |
| 4 | Germinal Center B Cell Undefined Human | 0.001085 | 50.74 | 346.39 |
| 5 | Marginal Zone B Cell Undefined Human | 0.001085 | 50.74 | 346.39 |
| 6 | Astrocyte Cortex Human | 0.001581 | 40.59 | 261.80 |
| 7 | Cholangiocyte Liver Human | 0.001581 | 40.59 | 261.80 |
| 8 | Neural Stem Cell Brain Human | 0.0002274 | 29.28 | 245.59 |
| 9 | Epithelial Cell Stomach Human | 0.002166 | 33.82 | 207.50 |
| 10 | Epithelial Cell Kidney Human | 0.002491 | 31.22 | 187.17 |

### Suppl. Table 4

#### A) Top upregulated signaling systems - GSEA Human Molecular Signatures Database (MSigDB)

| Index | Name | P-value | Odds Ratio | Combined score |
|---|---|---|---|---|
| 1 | Epithelial Mesenchymal Transition | 0.0005070 | 6.48 | 49.19 |
| 2 | p53 Pathway | 0.0005070 | 6.48 | 49.19 |
| 3 | Pancreas Beta Cells | 0.01706 | 10.67 | 43.42 |
| 4 | IL-2/STAT5 Signaling | 0.003253 | 5.35 | 30.62 |
| 5 | TGF-beta Signaling | 0.02993 | 7.79 | 27.33 |
| 6 | Coagulation | 0.03200 | 4.53 | 15.59 |

#### B) Top upregulated biological processes - Gene Ontology (GO) Molecular Function

| Index | Name | P-value | Odds Ratio | Combined score |
|---|---|---|---|---|
| 1 | Aldehyde Dehydrogenase (NAD+) Activity (GO:0004029) | 0.002166 | 33.82 | 207.50 |
| 2 | BMP Receptor Activity (GO:0098821) | 0.02475 | 50.24 | 185.84 |
| 3 | Pyruvate Transmembrane Transporter Activity (GO:0050833) | 0.02475 | 50.24 | 185.84 |
| 4 | Pyruvate Transmembrane Transporter Activity (GO:0050833) | 0.02475 | 50.24 | 185.84 |
| 5 | Secondary Active Monocarboxylate Transmembrane Transporter Activity (GO:0015355) | 0.02475 | 50.24 | 185.84 |
| 6 | Sphingosine N-acyltransferase Activity (GO:0050291) | 0.02963 | 40.19 | 141.43 |
| 7 | G Protein-Coupled Acetylcholine Receptor Activity (GO:0016907) | 0.02963 | 40.19 | 141.43 |
| 8 | G Protein-Coupled Neurotransmitter Receptor Activity (GO:0099528) | 0.02963 | 40.19 | 141.43 |
| 9 | Carbohydrate:Proton Symporter Activity (GO:0005351) | 0.02963 | 40.19 | 141.43 |
| 10 | MHC Class II Protein Binding (GO:0042289) | 0.02963 | 40.19 | 141.43 |

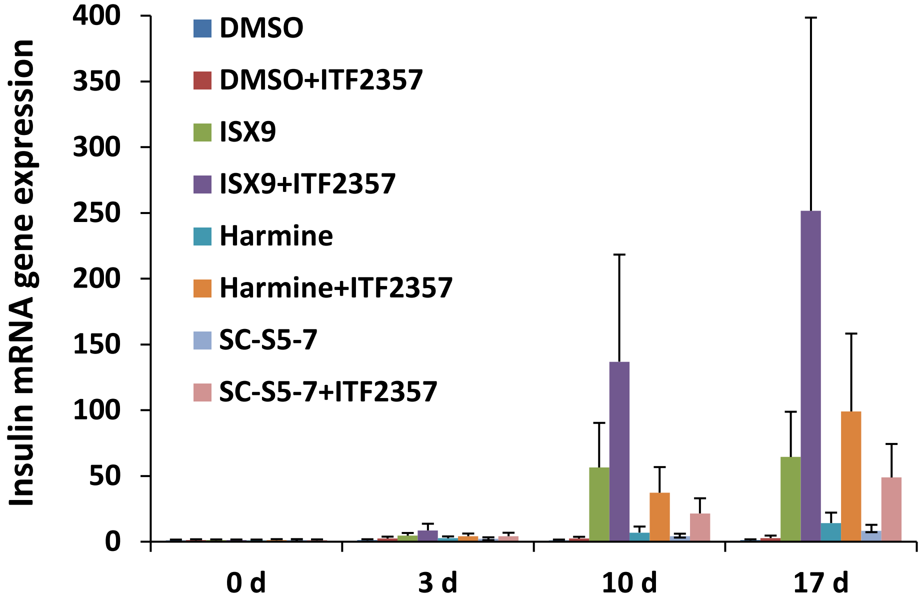


**Suppl. Fig 1. Comparison of treatments to induce insulin gene expression in islet organoids.** IPC clusters were treated with DMSO control, ISX9, Harmine, and differentiation medium SC-S5-7 alone and in combination with ITF2357 for 0 d, 3 d, 10 d, and 17 d
